# Increased abundance of nuclear HDAC4 impairs neuronal development and long-term memory

**DOI:** 10.1101/2021.02.04.429836

**Authors:** Patrick Main, Wei Jun Tan, David Wheeler, Helen L Fitzsimons

## Abstract

Dysregulation of the histone deacetylase HDAC4 is associated with both neurodevelopmental and neurodegenerative disorders, and a feature common to many of these disorders is impaired cognitive function. HDAC4 shuttles between the nucleus and cytoplasm in both vertebrates and invertebrates and alterations in the amounts of nuclear and/or cytoplasmic HDAC4 have been implicated in these diseases. In *Drosophila*, HDAC4 also plays a critical role in the regulation of memory however the mechanisms through which it acts are unknown. Nuclear and cytoplasmically-restricted HDAC4 mutants were expressed in the *Drosophila* brain to investigate a mechanistic link between HDAC4 subcellular distribution, transcriptional changes and neuronal dysfunction. Deficits in mushroom body morphogenesis, eye development and long-term memory correlated with increased abundance of nuclear HDAC4 but were associated with minimal transcriptional changes. Although HDAC4 sequesters MEF2 into punctate foci within neuronal nuclei, no alteration in MEF2 activity was observed on overexpression of *HDAC4*, and knockdown of *MEF2* had no impact on long-term memory, indicating that HDAC4 is likely not acting through MEF2. Similarly, deletion of the MEF2 binding site also had no impact on HDAC4-induced impairments in eye development, however it did significantly reduce the mushroom body deficits, thus nuclear HDAC4 acts through MEF2 to disrupt mushroom body development. These data provide insight into the mechanisms through which dysregulation of HDAC4 subcellular distribution impairs neurological function and provides new avenues for further investigation.

## Introduction

Dysregulation of the histone deacetylase HDAC4 results in impairments in neuronal development and formation of long-term memories in the *Drosophila* brain (Fitzsimons et al., 2013). Similarly in vertebrates, altered expression or subcellular distribution of HDAC4 is associated with both neurodevelopmental and neurodegenerative disorders, including 2q37 deletion syndrome (previously called brachydactyly mental retardation syndrome) (Williams et al., 2010;Morris et al., 2012;Villavicencio-Lorini et al., 2013), CDKL5 disorder (Trazzi et al., 2016), Alzheimer’s disease (Cao et al., 2008;Herrup et al., 2013;Shen et al., 2016), autism (Williams et al., 2010;Pinto et al., 2014) Huntington’s disease (Mielcarek et al., 2013a) and ataxia telangiectasia (Li et al., 2012;Herrup et al., 2013;Shen et al., 2016). Impaired cognitive function is a feature common to all of these disorders. HDAC4 shuttles between the nucleus and cytoplasm in both vertebrates and invertebrates (Chawla et al., 2003;Fitzsimons et al., 2013) and alterations in the amounts of nuclear and/or cytoplasmic HDAC4 have been implicated in these diseases (Cao et al., 2008;Sando et al., 2012;Herrup et al., 2013;Mielcarek et al., 2013a;Mielcarek et al., 2015;Shen et al., 2016). A frameshift mutation that results in nuclear accumulation of HDAC4 also results in features consistent with 2q37 deletion syndrome (Williams et al., 2010) and expression of the corresponding mouse variant of HDAC4 in the mouse brain causes deficits in spatial learning and memory (Sando et al., 2012). This may be initially surprising given that this mutant lacks a deacetylase domain, however HDAC4 is catalytically inactive in vertebrates and there are no global changes in histone acetylation resulting from knock out of HDAC4 in mice (Mielcarek et al., 2013b). Nuclear accumulation of HDAC4 has been observed in human Alzheimer’s disease post-mortem brains as well as the brains of Alzheimer’s mice with the abundance of nuclear HDAC4 correlating with increased clinical dementia scores in humans (Herrup et al., 2013;Shen et al., 2016), indicating that excess abundance of nuclear HDAC4 impairs cognitive function.

Nuclear import of HDAC4 requires a nuclear localization signal as well as binding of the transcription factor MEF2. Mutation of leucine 175 to alanine (L175A) within the MEF2 binding region of HDAC4 prevents MEF2 binding and results in cytoplasmic accumulation of HDAC4 (Wang and Yang, 2001). Nuclear export is regulated via activity-dependent CaM kinase-mediated phosphorylation of conserved serine residues S246, S467 and S623, which allows binding of the chaperone 14-3-3ζ (Grozinger and Schreiber, 2000;McKinsey et al., 2000;Wang et al., 2000;Wang and Yang, 2001;Zhao et al., 2001), and export from the nucleus (McKinsey et al., 2001). Mutation of these residues to alanine (3SA) results in nuclear retention (Chawla et al., 2003;Sando et al., 2012;Schlumm et al., 2013). In the nucleus, HDAC4 has been identified as a transcriptional regulator that represses transcription factors such as MEF2 (Miska et al., 1999). MEF2 family members promote neuronal survival and regulate both memory formation and dendrite morphogenesis (Flavell et al., 2006;Shalizi et al., 2006;Cole et al., 2012). Furthermore, they regulate multiple synapse-associated genes, and promote the development of inhibitory synapses while repressing excitatory synapse development (Barbosa et al., 2008).

*Drosophila* HDAC4 (DmHDAC4) shares 57% amino acid identity and 84% similarity to human HDAC4 (hHDAC4) across the deacetylase domain-containing C terminus, and 35% identity and 59% similarity across the whole protein and it is the sole Class IIa HDAC in *Drosophila*. The N-terminal elements critical for function and regulation of subcellular localization are conserved in DmHDAC4, including the MEF2 binding motif, serine residues for 14-3-3 mediated nuclear export, and a putative nuclear import signal (Fitzsimons et al., 2013). We previously found that overexpression of wild-type *HDAC4* in the adult mushroom body, a critical structure for formation of associative memory in *Drosophila* (McBride et al., 1999) resulted in impaired long-term memory (LTM) in a *Drosophila* model of courtship memory. The LTM deficits that we observed were specific to LTM as short-term memory and courtship activity were unaffected (Fitzsimons et al., 2013). This suggested that HDAC4 facilitates altered expression of genes that are required for LTM, however RNA-seq on *Drosophila* heads overexpressing *HDAC4* in the brain revealed very few changes (Schwartz et al., 2016). Moreover, it is likely that these effects are deacetylase independent (*Drosophila* HDAC4 is catalytically active) as overexpression of an *HDAC4* variant with a mutated H968 residue that disrupts deacetylase activity (Miska et al., 1999) also caused severe impairment of LTM (Fitzsimons et al., 2013). In addition, knockdown of *HDAC4* in the mushroom body of adult flies caused a similar phenotype, with male flies unable to form long-term memories, supporting the hypothesis that HDAC4 is a repressor of memory when in excess but also plays an essential role in normal LTM. We also carried out a screen for genes that interact genetically with *HDAC4* in eye development, and identified genes encoding proteins both that interact with and/or regulate the actin cytoskeleton and influence neuronal growth, as well as transcriptional machinery and genes involved in SUMOylation (Schwartz et al., 2016). Despite these advances, the specific mechanisms through which HDAC4 acts are still unclear, and complicating our understanding is that nuclear accumulation of HDAC4 goes hand-in-hand with cytoplasmic depletion, making it unclear whether cytoplasmic HDAC4 is required for learning and memory or whether it is merely being sequestered outside of the nucleus.

Here we aimed to further investigate the role of HDAC4, specifically to determine a mechanistic link between HDAC4 subcellular distribution, transcriptional changes and neuronal dysfunction. Mutant variants of HDAC4 that were sequestered in the nucleus or cytoplasm were compared to wild-type *Drosophila* and human HDAC4 to untangle the separate roles of nuclear and cytoplasmic HDAC4 in development and memory, as well as directly comparing the activity of *Drosophila* and human HDAC4s. Both mammalian HDAC4 and DmHDAC4 have been demonstrated to repress memory in their respective models, however, until now their subcellular distribution and respective roles in neurons have not been directly compared.

## Results

### Characterization of expression and subcellular distribution in the mushroom body

To characterize the contribution of subcellular pools of HDAC4 to the previously observed impairments in neuronal function, we employed transgenic flies expressing the human *HDAC4* mutants that are sequestered in the nucleus or cytoplasm. Nuclear and cytoplasmically restricted human HDAC4 mutants have been previously characterized (Cohen et al., 2007;Chen and Cepko, 2009); *3SA* encodes a nucleus-sequestered variant of hHDAC4 with three serine to alanine mutations that prevent nuclear export, and *L175A* encodes a cytoplasm-sequestered variant that harbors a leucine to alanine mutation that prevents nuclear import.

Prior to functional analyses, the subcellular distribution of wild-type hHDAC4 and the mutants was compared to that of wild-type DmHDAC4. Expression was driven in the mushroom body with the *OK107-GAL4* driver (Connolly et al., 1996;Aso et al., 2009) and induced in adult flies via temperature-dependent manipulation of GAL80ts, which represses GAL4 at the permissive temperature of 18°C. Repression is relieved by raising the temperature to 30°C, thus allowing transgene expression (McGuire et al., 2004) (Figure 1A). In comparison to GFP which was distributed relatively evenly through the cell, DmHDAC4 was distributed in both the axons and cell bodies of Kenyon cells with robust staining in the axons and in cytoplasmic haloes surrounding the nuclei (Figure 1B). Punctate foci were observed in a subset of nuclei as previously observed (Fitzsimons et al., 2013). Human HDAC4 also localized strongly to the axons, however there was a much lower presence of hHDAC4 in nuclei, which is likely due to the nuclear export sequence in hHDAC4 that appears absent in DmHDAC4. The cytoplasmically-restricted L175A mutant was completely absent from nuclei, as expected. In contrast, neuronal nuclei of brains expressing 3SA contained numerous punctate foci, indicating nuclear retention, although it was not completely excluded from the cytoplasm as significant staining was also observed in the axons.

**Figure 1.**
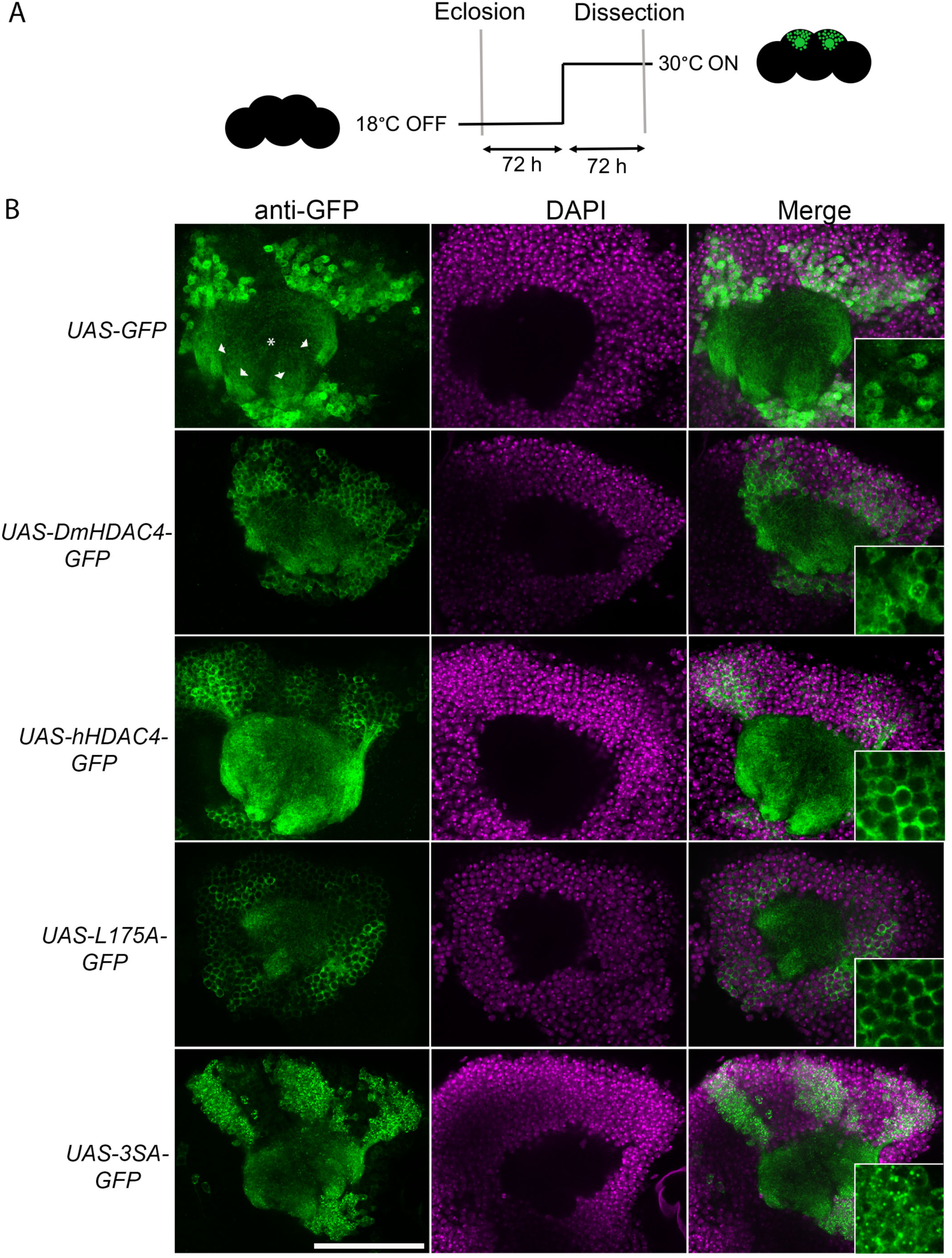
Expression and subcellular localization of GFP-tagged HDAC4 constructs in the mushroom body. A. Schematic diagram describing protocol for induction of expression in the adult mushroom body. All genotypes were generated by crossing *OK107-GAL4; tubP-GAL80*_*ts*_ females to males carrying the indicated UAS-transgene. All *HDAC4* transgenes were fused in frame to GFP. Flies were raised at 18°C, at which temperature GAL80 represses GAL4, until after eclosion when the temperature was raised to 30°C. At this temperature GAL80 is inactivated, allowing GAL4 to induce transgene expression. Brains were dissected after 72 hours at 30°C. B. Whole-mount brains were subjected to immunohistochemistry with anti-GFP (green) and counterstained with DAPI (magenta). Optical sections (1 μm) through the calyx of the mushroom body are shown. Asterisks label the calyx and arrowheads indicate the four bundles of Kenyon cell axons that originate in the calyx and project anteriodorsally to form the lobes. Scale bar = 40 μm. Insets in the right panels are a 4x magnification of cell bodies.

### Increased nuclear HDAC4 impairs neuronal development

We next assessed the impact of increasing either nuclear or cytoplasmic HDAC4 on mushroom body development as we had previously observed that overexpression of wild-type *DmHDAC4* disrupts normal axon branching, elongation and termination (unpublished data). The mushroom body is comprised of three classes of intrinsic neurons, the α/β, α’/β’ and γ Kenyon cells. The axons of the three Kenyon cell subtypes are bundled together to form a ventrally projecting peduncle, which then splits to form lobes. The α/β and α’/β’ axons both bifurcate to form the vertical α and α’ lobes and the medial β and β’ lobes, while the γ neuron axons form a single medial lobe. The integrity of the mushroom body can be assessed by via visualization of the α, β and γ lobe marker FasII (Freymuth and Fitzsimons, 2017) (Figure 2). *elav-GAL4* driven pan-neuronal expression of wild-type *DmHDAC4* in post-mitotic neurons disrupted normal development in 95% of brains, manifesting as impairments in α/β lobe elongation and β-lobe fusion, which occurs when axons erroneously cross the midline. Defects resulting from expression of *hHDAC4* were significantly less pronounced than *DmHDAC4*, with 48% penetrance and the majority of the defects were thinner lobes (Table 1). Expression of *3SA* severely disrupted development with all brains displaying structural abnormalities, whereas in contrast, 85% of brains expressing cytoplasmic *L175A* appeared wild-type.

**Table 1.** Frequency of mushroom body phenotypes resulting from expression of *HDAC4* mutants. The percentage of brains displaying each phenotype was calculated from the total number of brains analyzed for each genotype (n). Statistical analysis was performed with Fisher’s Exact Test. Expression of *DmHDAC4* resulted in significantly more abnormal brains than *hHDAC4* (p=0.0007), as did *3SA* (p<0.00001). *3SA* expression disrupted development to a significantly greater degree than *L175A* (p<0.00001), which was not significantly different to the *w(CS10)* control (p=0.104)

| Genotype | *elav>+* | *elav> DmHDAC4* | *elav>hHDAC4* | *elav>3SA* | *elav>L175A* |
| --- | --- | --- | --- | --- | --- |
| *n* | 23 | 21 | 23 | 31 | 21 |
| No defects | 100% | 5% | 52% | 0% | 85% |
| Thin $\alpha$ or $\beta$ lobes | 0% | 19% | 30% | 19% | 10% |
| Fused $\beta$ lobes | 0% | 14% | 9% | 3% | 5% |
| Thin $\alpha$ or $\beta$ lobes and fused $\beta$ lobes | 0% | 52% | 4% | 45% | 0% |
| Absent $\alpha$ or $\beta$ lobe | 0% | 10% | 4% | 23% | 0% |
| Multiple absent lobes | 0% | 0% | 0% | 6% | 0% |

**Figure 2.**
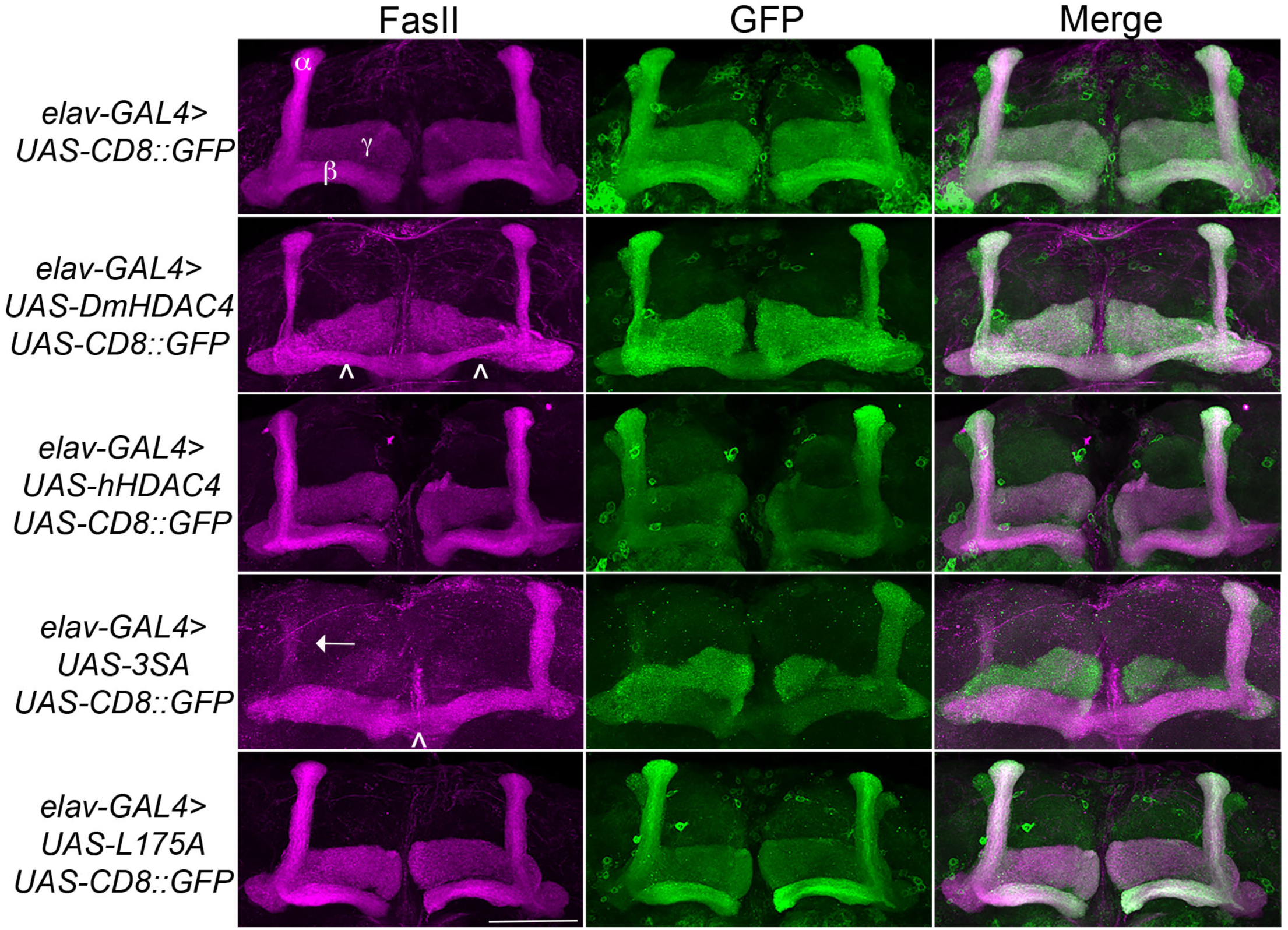
Nuclear accumulation of HDAC4 disrupts mushroom body development. Immunohistochemistry with anti-FasII and anti-GFP antibodies on whole mount brains expressing FLAG-tagged HDAC4 variants and CD8::GFP as a counterstain to visualize all the lobes. All images are frontal confocal projections through the mushroom body region of the brain. All genotypes were generated by crossing *elav-GAL4; UAS-CD8::GFP* females to males carrying each *UAS-HDAC4* transgene and to the *w(CS10)* control. Brains displaying representative phenotypes are shown. Scale bar = 50 μm. α, β and γ lobes of the mushroom body are labeled in white. Thin and fused β lobes (^) are evident in a brain expressing *Drosophila HDAC4*. A brain expressing human *HDAC4* appears normal. Missing α lobe (arrow) and fused β lobes (^) are evident in a brain expressing *3SA*. A brain expressing *L175A* appears normal.

We also examined whether nuclear accumulation of HDAC4 impacted eye development. The *Drosophila* eye is rich in neuronal photoreceptors and defects in its structure are readily observable, making it an excellent model for genetic analysis of neuronal pathways. We previously showed that expression of *HDAC4* in the post-mitotic eye via the glass multimer reporter (*GMR*) driver (Freeman, 1996) causes a disruption of the hexagonal ommatidia and bristles between them as well as a reduction in pigmentation, which was rescued via RNAi-mediated knockdown of *HDAC4* (Schwartz et al., 2016). Here we found that expression of *3SA* resulted in more severe deficits than *DmHDAC4* with reduced pigmentation, fused ommatidia and disorganized bristles, which was not observed with wild-type *hHDAC4* nor *L175A* (Figure 3).

**Figure 3.**
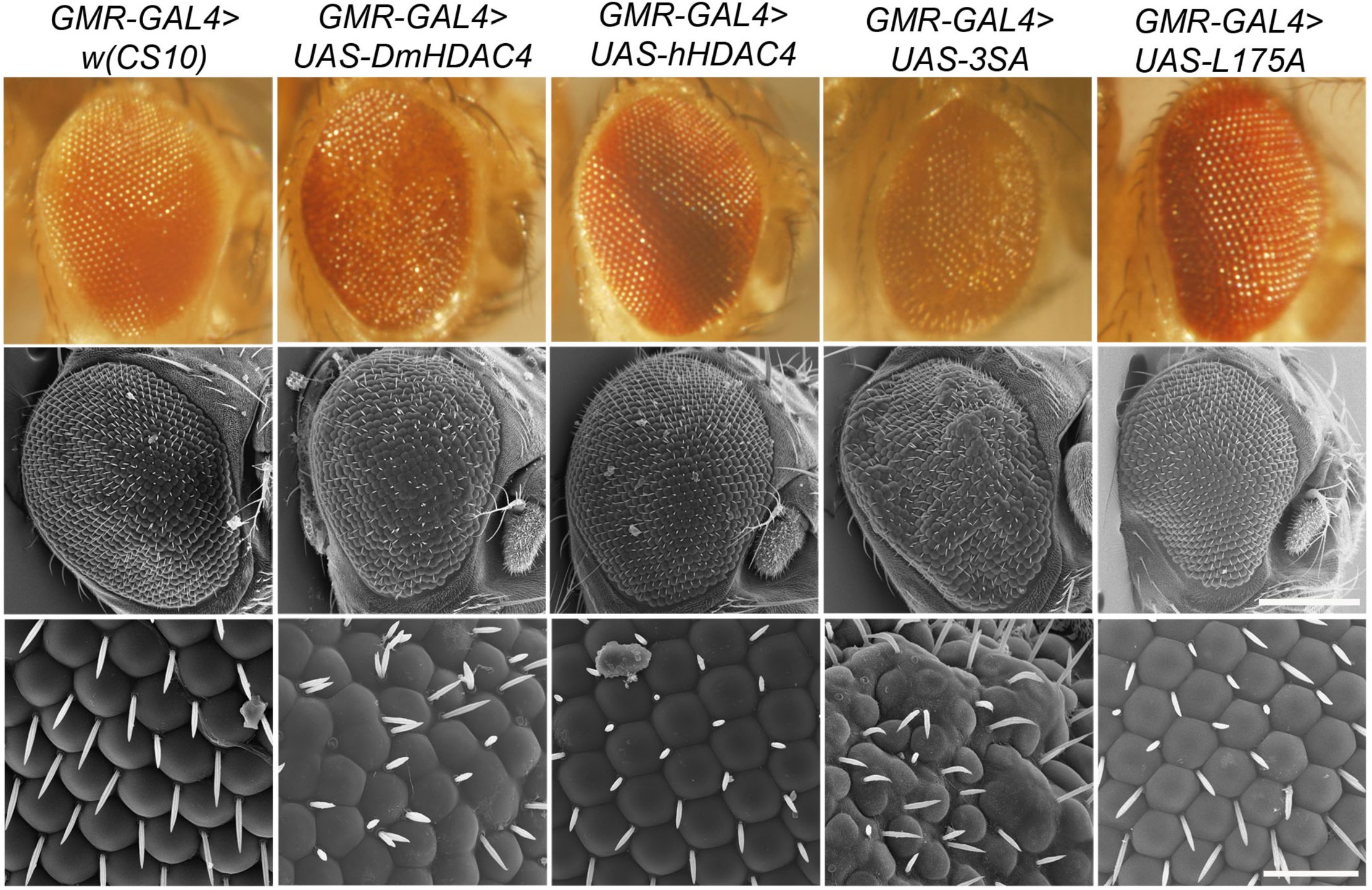
Nuclear accumulation of HDAC4 disrupts eye development. Stereomicrographs and scanning electron micrographs of *Drosophila* eyes expressing FLAG-tagged *HDAC4* variants. Genotypes were generated by crossing *GMR-GAL4* females to males carrying each *UAS-HDAC4* transgene and to the *w(CS10)* control. Top panel: Stereomicrographs, 110x magnification. Middle panel: SEM, 250x magnification. Scale bar = 200 μm. Lower Panel: SEM, 1500x magnification SEM. Scale bar = 30 μm.

### Increased nuclear HDAC4 impairs long-term memory

We next investigated whether nuclear HDAC4 was responsible for the long-term memory deficits we have previously observed on overexpression of *DmHDAC4* in the adult mushroom body (Fitzsimons et al., 2013). *OK107-GAL4*-driven expression was induced in the adult mushroom body (Figure 4A), and the courtship suppression assay was then used to evaluate memory. In this assay, a male fly’s courtship behavior is modified by its previous experience of rejection by a mated female, providing a quantitative assessment of associative memory (Figure 4B). Expression of *DmHDAC4* impaired LTM (Figure 4C) and while flies expressing *hHDAC4* displayed reduced LTM, this was not significant. Comparison of *hHDAC4* to *L175A* and *3SA* revealed that expression of *3SA* impaired LTM to a significant level whereas *L175A* did not (Figure 4C). This was not due to a reduction in courtship behavior as naive flies of each genotype displayed normal courtship (Figure 4D). Taken together these data indicate that the nuclear accumulation of HDAC4 correlates with structural deficits in the developing mushroom body and eye, and impairs formation of LTM when expressed in the adult mushroom body.

**Figure 4.**
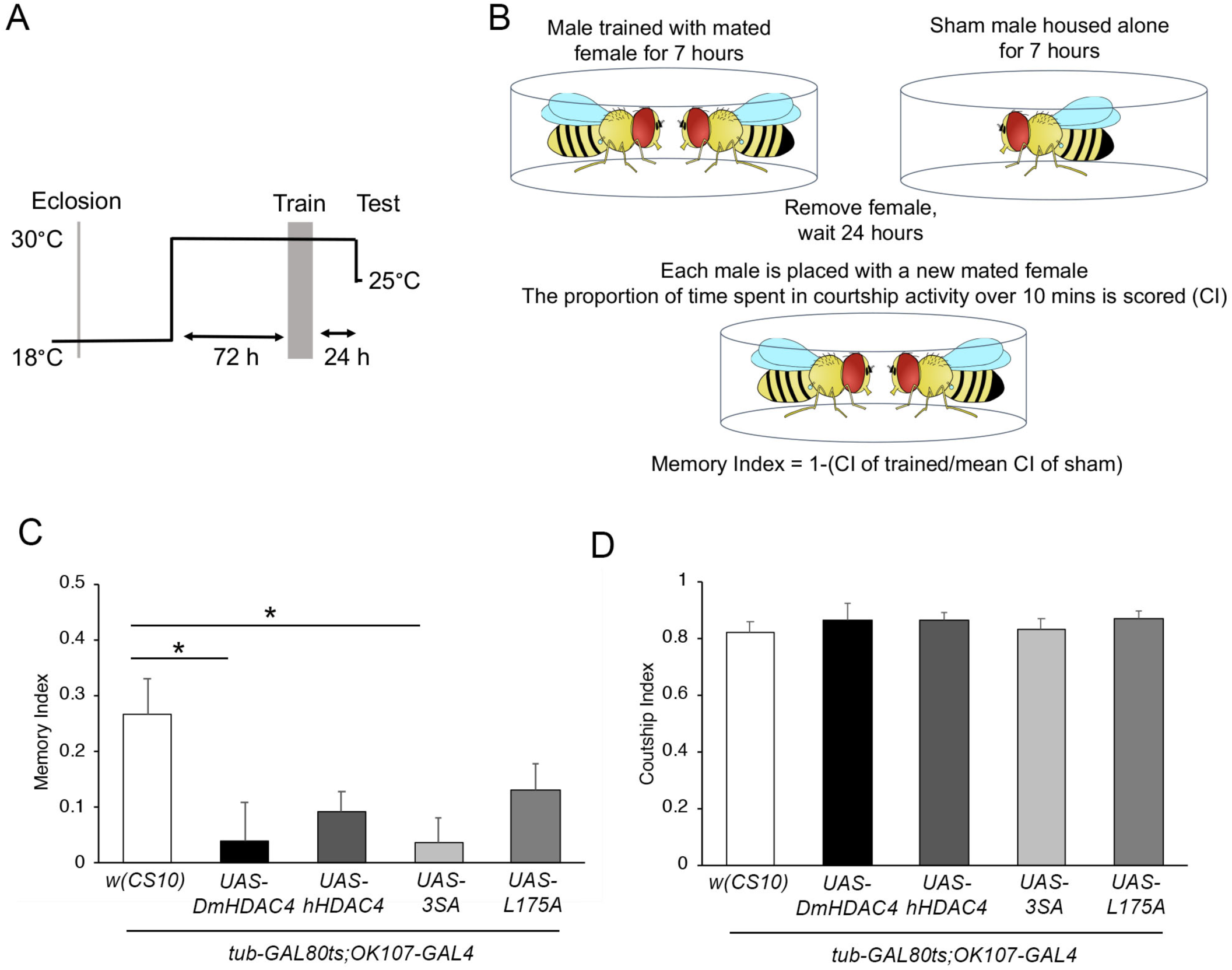
Nuclear accumulation of HDAC4 disrupts long-term memory. A. Schematic diagram describing the protocol for induction of expression in the adult mushroom body. All genotypes were generated by crossing *OK107-GAL4; tubP-GAL80*_*ts*_ females to males carrying the indicated UAS-transgene or to the *w(CS10)* control. Flies were raised at 18°C until after eclosion, then placed at 30°C to induce expression in the adult brain. They were trained 72 hours later and remained at 30°C until 1 hour prior to testing at which time they were equilibrated to the testing temperature of 25°C. B. Schematic diagram describing the courtship suppression assay. C. 24-hour courtship LTM was significantly impaired by expression of *DmHDAC4* and *3SA* in the adult brain (ANOVA, F_(4,209)_=3.59, p<0.007; post-hoc Tukey’s HSD, ^*^p<0.05). D. Courtship activity of untrained flies was not altered by expression of any of the HDAC4 variants (ANOVA, F_(4,214)=_0.45, p=0.772).

### Characterization of transcriptional roles of HDAC4

We previously demonstrated that increased nuclear abundance of *Drosophila* HDAC4 results in the transcription factor MEF2 being sequestered into HDAC4-positive punctate nuclear foci (Fitzsimons et al., 2013). Similarly, expression of *3SA* also causes a redistribution of MEF2 from a relatively uniform distribution into numerous puncta in the nucleus, confirming that hHDAC4 also interacts with DmMEF2 (Figure 5A). HDAC4 has been proposed to act as an E3 ligase, as it was demonstrated to enhance SUMOylation of MEF2 in mammalian cells (Gregoire and Yang, 2005;Zhao et al., 2005). We previously showed that *HDAC4* interacts genetically with the SUMOylation machinery and that the SUMO E3 ligase Ubc9 is required for LTM in *Drosophila*, but increased nuclear HDAC4 did not appear to alter global SUMOylation, nor facilitate SUMOylation of candidates MEF2, CREB, CaMKII or itself (Schwartz et al., 2016).This does not preclude that there could be changes in SUMOylation of specific unknown targets, and we hypothesized that HDAC4 may sequester SUMOylated proteins into the nuclear puncta observed in 3SA-expressing nuclei. However we saw no obvious colocalization of SUMO with 3SA-positive foci (Figure 5B).

**Figure 5.**
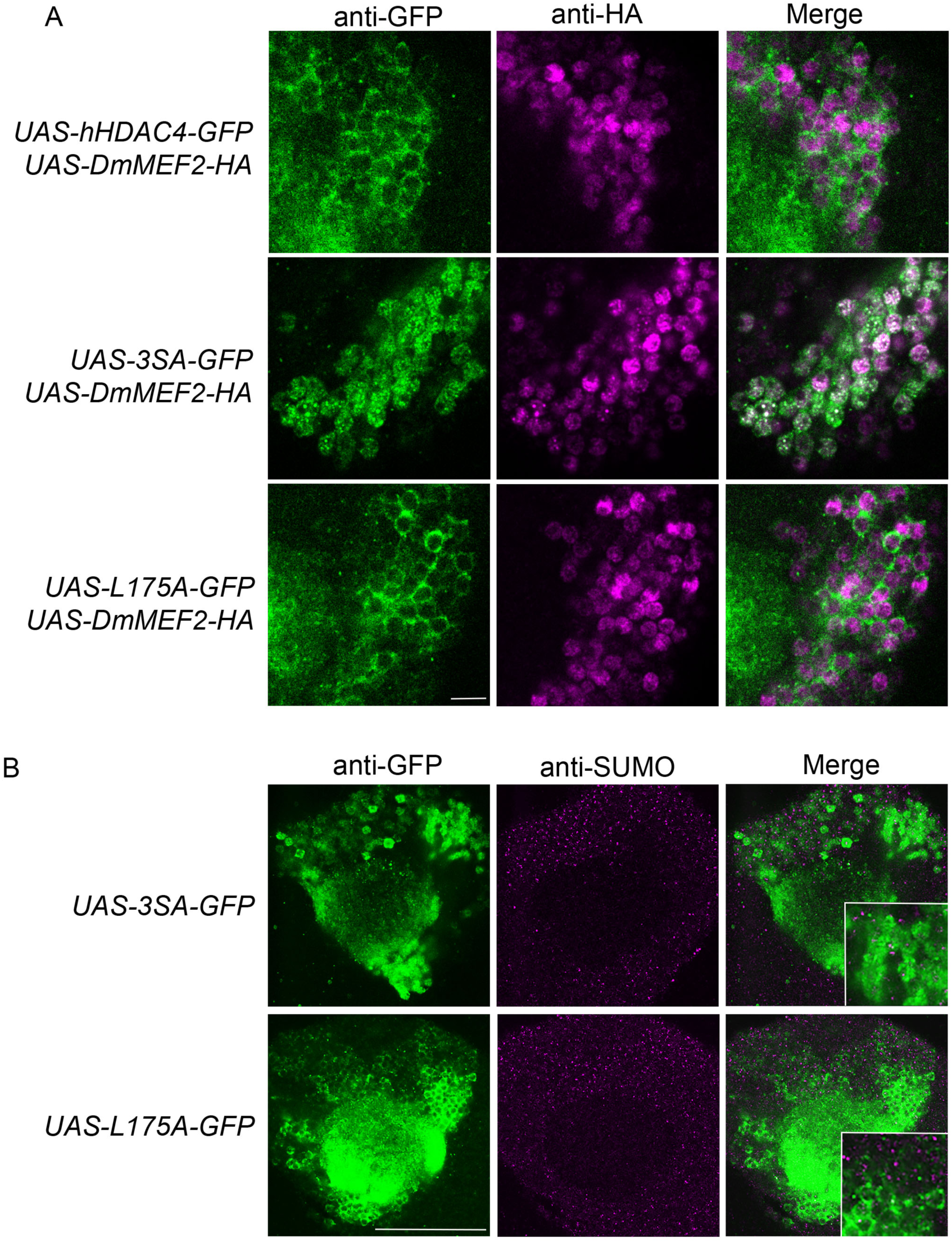
3SA distributes with MEF2 but not SUMO in nuclei. Immunohistochemistry on whole mount brains expressing the indicated transgenes. All images are 0.5 μm optical sections through the Kenyon cells at the level of the calyx. All genotypes were generated by crossing *tubP-GAL80ts; OK107-GAL4* females to males of the indicated genotype. Expression was induced in the adult brain by raising the flies at 18°C and placing at 30°C 72 hours prior to dissection. Representative images from each genotype are shown. A. 3SA co-distributes with MEF2 in nuclei. Whole-mount brains were subjected to immunohistochemistry with anti-HA (magenta) and anti-GFP (green). Scale bar = 4 μm. B. 3SA does not co-distribute with SUMO in nuclei. Whole-mount brains were subjected to immunohistochemistry with anti-SUMO (magenta) and anti-GFP (green). Optical sections (1 μm) through the Kenyon cell bodies are shown. Scale bar = 40 μm. Insets show 2x magnification of Kenyon cell bodies.

*DmHDAC4* and *MEF2* have been shown to interact genetically in a rough eye screen, indicating they act in the same molecular pathway during eye development (Schwartz et al., 2016) and MEF2 is required for normal mushroom body development in *Drosophila* (Crittenden et al., 2018). Together these data suggest the possibility that repression of MEF2-induced transcription may be a mechanism through which HDAC4 impairs memory and brain development in *Drosophila*. In this case, reduction of *MEF2* expression should be expected to impair LTM, however RNAi knockdown of *MEF2* had no significant impact on LTM (Figure 6A). Interestingly, overexpression of *MEF2* did impair LTM (Figure 6B), suggesting it is acting as a memory repressor. Neither knockdown nor overexpression of *MEF2* altered courtship activity itself (Figure 6C). Knockdown and overexpression of *MEF2* was verified via RT-qPCR and immunohistochemistry, respectively (Figure S1).

**Figure 6.**
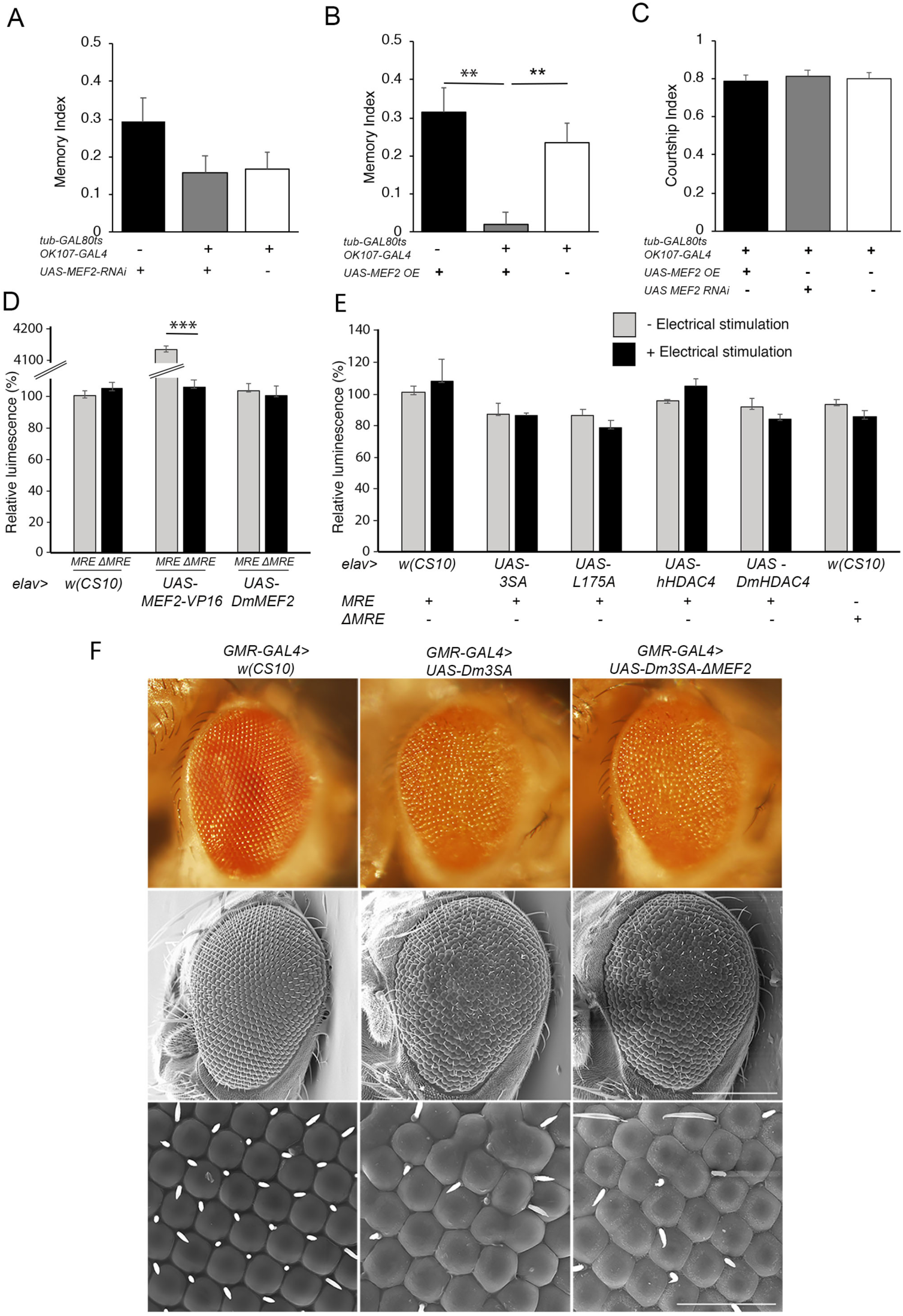
Functional analysis of the MEF2/HDAC4 interaction. A-C. All genotypes were generated by crossing *OK107-GAL4; tubP-GAL80*_ts_ females to males of the indicated genotype. Expression was induced in the adult brain by raising the flies at 18°C then placing at 30°C for 72 hours. The courtship suppression assay was carried out as per Fig 4A. A. *MEF2* knockdown in Kenyon cells does not alter LTM (ANOVA, F_(2,158)=_2.06, p=0.13). B. Overexpression of *MEF2* in Kenyon cells impairs LTM (ANOVA, F_(2,156)_=10.91, p<0.0001; post-hoc Tukey’s HSD,^*^p<0.01). C. Courtship activity was not altered by knockdown or overexpression of *MEF2* (ANOVA, F_(2,174)=_1.62 p=0.201). D. *elav-GAL4* driven expression of *MEF2-VP16* activated expression of *MRE-luc* but not *ΔMRE-luc* ANOVA, F_(5,18)_=1645, p<0.000001; post-hoc Tukey’s HSD, ^*^p<0.000001). Expression of *DmMEF2* did not increase luciferase activity above background levels in *MRE-luc* brains. E. Electrical stimulation did not increase luciferase above background levels (*w(CS10*) control +/-electrical stimulation, *t*-test t_(5)_=0.691, p=0.520) and there was no alteration in luciferase activity on expression of wild-type or mutant *HDAC4* (ANOVA F_(5,17)_ =1.46, p=0.253). F. Stereomicrographs and scanning electron micrographs of *Drosophila* eyes in which the indicated transgenes are expressed with the *GMR-GAL4* driver. Top panel: Stereomicrographs. Middle panel: SEM 250x magnification. Scale bar = 200 μm. Lower Panel: SEM, 1500x magnification. Scale bar = 40 μm.

We further investigated the interaction between HDAC4 and MEF2 *in vivo* with flies carrying a MEF2-response element (MRE) fused to luciferase. We confirmed that this cassette was functional *in vivo* by co-expression of a constitutively active *MEF2* transgene which consists of the MEF2 DNA binding domain fused to the VP16 activation domain (Chawla et al., 2003). When driven by *elav-GAL4*, MEF2-VP16 activated expression of luciferase in the adult brain, whereas no induction was observed in flies carrying ΔMRE-luc confirming that the system was appropriately responsive to MEF2 *in vivo* (Figure 6D). Despite this, there was no significant difference in basal luciferase activity between MRE and ΔMRE flies, suggesting minimal endogenous MEF2 activity in the wild-type brain under basal conditions, and furthermore luciferase activity was not induced on overexpression of *Drosophila MEF2*. As MEF2 is activated by synaptic activity in mammals (Flavell et al., 2006) we induced neuronal depolarization via electrical stimulation of the brain with standard parameters used for olfactory conditioning (Qiu and Davis, 1993). However, this also resulted in no significant increase in luciferase activity above basal levels (Figure 6E). Following our finding that overexpression of *MEF2* impaired memory, we investigated whether nuclear HDAC4 could possibly facilitate an increase in MEF2 activity, however co-expression of wild-type or mutant *HDAC4* had no significant effect on luciferase activity in the absence of electrical stimulation, and there was no significant change in luciferase activity for any transgene when electrical stimulation was applied (Figure 6E).

To further examine the nature of a relationship between HDAC4 and MEF2, we generated flies carrying *3SA* with a mutated MEF2 binding site to determine whether the *3SA* phenotype was dependent on MEF2 binding. We firstly generated the corresponding *3SA* mutation in *DmHDAC4* to confirm conserved functionality between *Drosophila* and human HDAC4 and observed that like human *3SA, Drosophila 3SA* expression in the eye resulted in loss of pigmentation, fusion of ommatidia and missing or misplaced bristles (Figure 6F). Mutation of the MEF2 binding site (*Dm3SA-ΔMEF2*) resulted in a phenotype indistinguishable from *Drosophila 3SA* alone, indicating that an interaction with MEF2 is not required for the nuclear HDAC4-induced phenotype. In contrast to the eye phenotype, when examined for the effects on axon elongation and termination in the mushroom body, expression of *Drosophila 3SA* resulted in a similar phenotype to human *3SA* with missing and fused lobes, but this phenotype was reduced to close to wild-type in brains expressing *Dm3SA-ΔMEF2* (Table 2), indicating that MEF2 binding is important for the 3SA-induced developmental defects in the mushroom body.

**Table 2.** Frequency of mushroom body phenotypes resulting from expression of *Dm3SA* in the presence and absence of the MEF2 binding site. The percentage of brains displaying each phenotype was calculated from the total number of brains analyzed for each genotype (n). Statistical analysis was performed with Fisher’s Exact Test. Expression of *Dm3SA* resulted in significantly more abnormal brains than *Dm3SAΔMEF2* (p<0.00001).

| Genotype | *elav>+* | *elav> Dm3SA* | *elav>Dm3SA+ΔMEF2* |
| --- | --- | --- | --- |
| *n* | 23 | 25 | 21 |
| No defects | 100% | 4% | 71% |
| Thin $\alpha$ or $\beta$ lobes | 0% | 0% | 14% |
| Fused $\beta$ lobes | 0% | 40% | 5% |
| Thin $\alpha$ or $\beta$ lobes and fused $\beta$ lobes | 0% | 52% | 0% |
| Absent $\alpha$ or $\beta$ lobe | 0% | 0% | 10% |
| Multiple absent lobes | 0% | 4% | 0% |

Given that we could not detect transcriptional activity of MEF2, we examined whether expression of *3SA* resulted in any changes in gene expression via RNA-seq. Previous RNA-seq on heads of flies overexpressing *DmHDAC4* in the adult fly brain revealed few significant transcriptional changes and we reasoned that this may be due to a dilution effect, given that wild-type DmHDAC4 is nuclear in only a subset of cells. We therefore performed RNA-seq on heads of flies expressing *3SA* in the adult brain. Developmental effects were avoided by raising flies at 18°C and inducing expression via temperature-dependent repression of GAL80ts at 30°C following eclosion of adults. Very few transcriptional changes were observed (Figure 7A, Table S1), leading us to conclude that even when sequestered in all neuronal nuclei, HDAC4 elicits only small changes in transcription in the fly brain.

**Figure 7.**
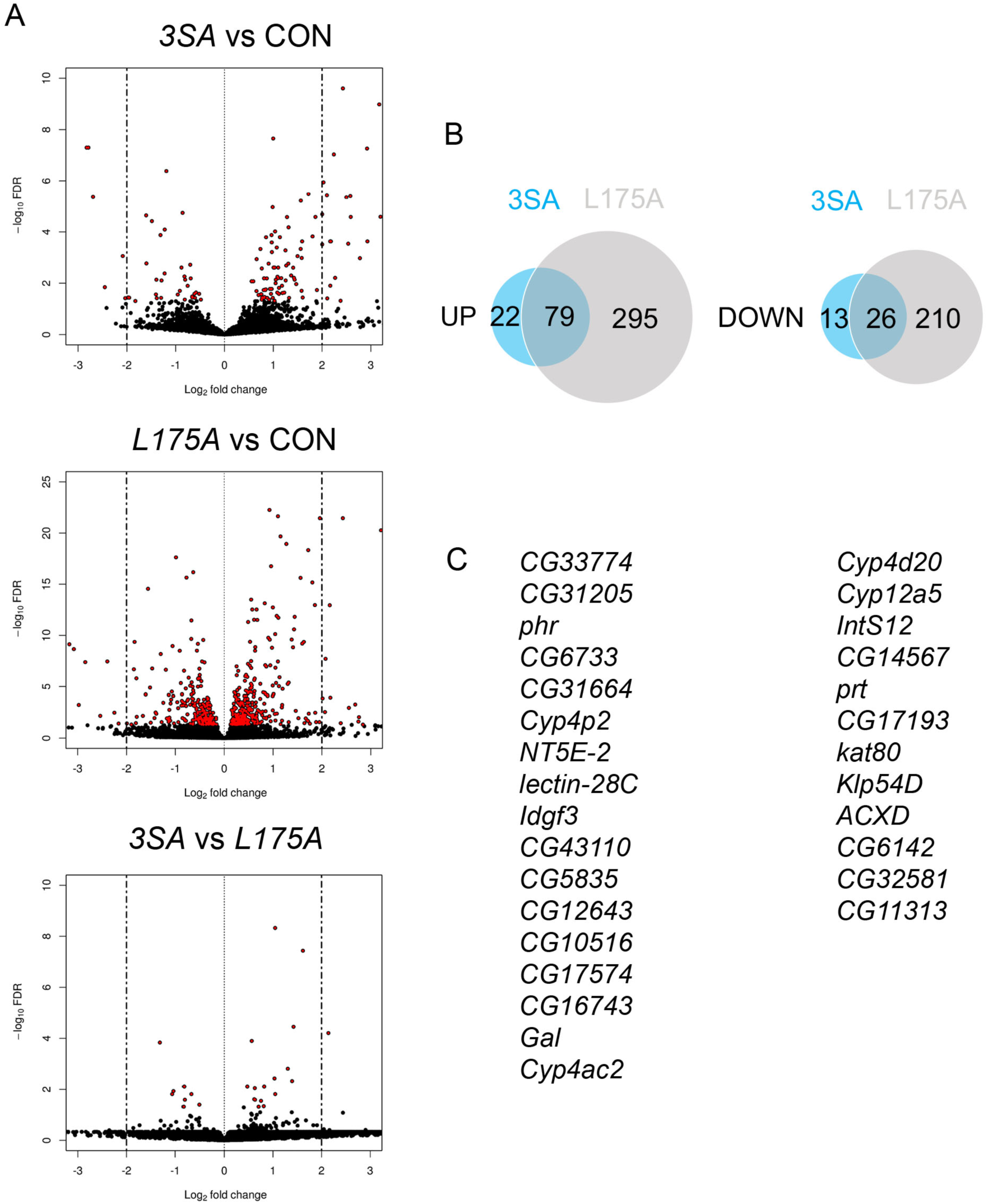
Transcriptional changes resulting from expression of *3SA* and *L175A*. A. Genotypes were generated by crossing *tubP-GAL80ts; OK107-GAL4* females to *UAS-FLAG-3SA, UAS-FLAG-L175A* or *w(CS10)* males (indicated as CON). Expression was induced in the adult brain by raising the flies at 18°C and placing at 30°C 72 hours prior to dissection. Volcano plots indicate differential expression between treatment groups. The horizontal axes indicate the Log2-fold change in expression and the vertical axis indicates the negative Log10 of the false discovery rate. Genes with significant differential expression (i.e. FDR < 0.05) are highlighted in red. Expression of *L175A* resulted in a higher number of differentially expressed genes than *3SA* (2-sample test for equality of proportions (X-squared = 296.89, df = 1, p-value<0.01)). B. Venn diagrams indicating the number of differentially regulated genes common to both control vs *L175A* and control vs *3SA* data sets. C. List of genes significantly altered in expression by *3SA* but not *L175A*. Only 29 genes were common to both data sets.

Surprisingly, there were significantly more differentially expressed genes in the *L175A* group versus *3SA* and the majority of genes that were differentially regulated by *3SA* expression were also altered by *L175A* (Fig 7B). Given that L175A appeared entirely cytoplasmic but 3SA was not completely restricted to the nucleus with some distribution in the cytoplasm (Figure 1B), it is a logical assumption that the change in expression of these genes was mediated by cytoplasmic HDAC4. A direct comparison of *3SA* and *L175A* revealed only 29 genes were significantly differentially expressed with a False Discovery Rate < 0.05 (Fig 7C, Table S1) and only four of these displayed more than a Log2-fold change in expression. Gene ontology (GO) analysis using the differentially expressed list from the *L175A* and *3SA* comparison identified enrichment of only four molecular functions: monooxygenase activity, paired donor oxidoreductase activity, heme binding and iron ion binding; and the source of these GO terms was only four genes that encode cytochrome P450 superfamily proteins (Table S1).

## Discussion

Here we demonstrate that the neurological impairments resulting from increased *HDAC4* expression correlate with increased abundance of nuclear HDAC4. Expression of wild-type and cytoplasmic human *HDAC4* resulted in minimal phenotypic changes whereas nuclear accumulation resulted in severe defects in mushroom body and eye development, as well as disruption of long-term memory. These data establish that *Drosophila* is an informative model for investigation of the molecular mechanisms that underlie disorders associated with nuclear accumulation of HDAC4.

Expression of *3SA* in nuclei of post-mitotic neurons caused deficits in elongation and termination of Kenyon cell axons. In contrast, in the mouse forebrain, expression of a nuclear-restricted truncated mutant of *HDAC4* did not alter gross architecture of the hippocampus nor neuronal survival as evidenced by number of dentate granule cells and densities of their synapses (Sando et al., 2012). In addition, brain-specific knockout of *HDAC4* in the mouse has no major impact on normal morphology or survival (Price et al., 2013) and nor does double *HDAC4/HDAC5* knockout (Zhu et al., 2019) suggesting HDAC4 does not play a role in development in an otherwise normal rodent brain. However vertebrates have four Class IIa HDACs while HDAC4 is the sole Class IIa HDAC in *Drosophila*, which avoids effects of redundancy and allows the basic functions of Class IIa HDACs to be uncovered. There are some instances however, in which nuclear accumulation of HDAC4 impacts brain development in vertebrates when associated with existing disease. Cyclin dependent kinase-like 5 (CDLK5) disorder is a neurodevelopmental disorder associated with intellectual disability, and *CDLK5* mutant mice display deficits in neuronal survival and maturation as well as impaired hippocampal-dependent memory. HDAC4 is a phosphorylation target of CDLK5 and in its absence, nuclear entry of unphosphorylated HDAC4 is associated with reduced histone H3 acetylation and modest repression of MEF2. Treatment with an HDAC4 inhibitor restores memory and reverses defects in neuronal maturation and death, which is highly suggestive that the deficits associated with *CDLK5* disorder are a result of nuclear accumulation of HDAC4 (Trazzi et al., 2016).

The expression of *3SA* in the adult mushroom body also abolished the formation of long-term memory, which is consistent with findings that a truncated mutant of human HDAC4 that accumulates in nuclei impairs memory (Sando et al., 2012). It is notable that this mutant also lacks a deacetylase domain, suggesting a deacetylase-independent mechanism. Taken together, it is evident that when overabundant in the nucleus, HDAC4 disrupts memory in both vertebrates and invertebrates. We previously identified that *HDAC4* interacts genetically with several cytoskeletal regulators including *Moesin* and *Ankyrin2*, although it does not regulate their expression (Schwartz et al., 2016). While the nature of the interaction is yet to be determined, we also showed that knockdown of *Moesin* resulted in similar mushroom body deficits and impaired long-term memory (Freymuth and Fitzsimons, 2017) as does knockdown of *Ankyrin2* (unpublished data). Together these data suggest that nuclear HDAC4 may be an upstream regulator of molecular pathways that regulate rearrangement of the actin cytoskeleton.

The most obvious mechanism through which HDAC4 would alter neuronal development and memory would be modulation of gene expression, however we previously found that overexpression of *DmHDAC4* was associated with only minor transcriptional changes (Schwartz et al., 2016). As the neuronal distribution of HDAC4 is largely cytoplasmic we reasoned that any significant changes in neuronal populations may have been diluted and could be uncovered by forcing HDAC4 to accumulate in nuclei of all neurons. However again we saw minimal changes resulting from forced nuclear accumulation via expression of *3SA* and surprisingly the majority of differential gene expression was a result of cytoplasmic accumulation of HDAC4. Comparison with our previous RNA-seq data in which *DmHDAC4* was overexpressed (Schwartz et al., 2016) revealed that of the 26 genes (not including *w* and *DmHDAC4*) differentially expressed in that study, 14 were also differentially expressed following expression of *L175A*, approximately 54% of the total. These data indicate conservation of function of HDAC4 between humans and *Drosophila* and surprisingly, that the changes elicited by DmHDAC4 are very likely through its cytoplasmic presence. How HDAC4 facilitates alterations in gene expression from the cytoplasm is unknown, but it could be through sequestration of transcriptional regulators in the cytoplasm, which has previously been demonstrated for the transcription factor ATF4, whereby increased cytoplasmic HDAC4 redistributed ATF4 from the nucleus to the cytoplasm (Zhang et al., 2014). It is clear that in the data we present here, these transcriptional changes resulting from increased cytoplasmic HDAC4 are not associated with any observed neurological phenotypes. However knockdown of *HDAC4* impairs LTM in *Drosophila* (Fitzsimons et al., 2013) as does brain-specific knockout of *HDAC4* (Kim et al., 2012) and *HDAC4/5* (Zhu et al., 2019) in mice, indicating it plays an essential role in normal memory formation. It therefore warrants further investigation as to whether the same genes and/or pathways are altered in the opposite direction upon reduction of HDAC4, which would provide insight into whether loss of cytoplasmic HDAC4 contributes to the memory deficits resulting from a reduction in *HDAC4* expression.

We observed that both human and *Drosophila* HDAC4 interact with MEF2, and when abundant in the nucleus they sequester MEF2 into punctate nuclear foci in Kenyon cells. Given that Kenyon cells are critical to formation of memory, and that HDAC4 is well established as a repressor of MEF2, it was a logical assumption that HDAC4 may impair memory by repressing MEF2. However the lack of transcriptional changes does not support this and moreover, knock down of *MEF2* in the adult brain had no impact on LTM. On the contrary, overexpression of *MEF2* significantly impaired LTM. This is consistent with evidence that MEF2 is a memory repressor in mice; inhibitory phosphorylation of MEF2 in the murine hippocampus or nucleus accumbens is associated with the formation of normal spatial and fear memories and overexpression of MEF2 in the dentate gyrus and amygdala blocks both spatial memory and fear memory, respectively (Cole et al., 2012). Together these data provide evidence that the mechanism through which nuclear sequestration of HDAC4 impairs memory in *Drosophila* is likely not via MEF2 repression. We similarly demonstrated that in the developing eye, 3SA does not require MEF2 binding to impair development. The situation is different in the mushroom body, in which expression of *MEF2* is required for normal mushroom body development (Crittenden et al., 2018). When *3SA* was overexpressed, similar phenotypes were observed to those when *MEF2* was knocked down (unpublished observations), but mutation of the MEF2 binding site prevented most of those 3SA-induced developmental defects.

Here we have shown that the irreversible accumulation of HDAC4 in the nucleus is associated with neurological dysfunction, and both we and others have observed that when overexpressed, HDAC4 aggregates into punctate foci within nuclei (Miska et al., 1999;Borghi et al., 2001;McKinsey et al., 2001;Fischle et al., 2002;Fitzsimons et al., 2013). The propensity of HDAC4 to aggregate stems from its glutamine-rich N-terminus which forms an alpha helix that can assemble into tetramers and further oligomerize. Interestingly, HDAC4-positive nuclear inclusions have been associated with pathogenesis of neurological disorders including Lewy bodies in Parkinson’s disease and intra-nuclear inclusions in intra-nuclear inclusion disease (Takahashi-Fujigasaki and Fujigasaki, 2006). These inclusions also contain SUMO (Takahashi-Fujigasaki et al., 2006), which was of interest given our previous finding that *HDAC4* interacts in the same molecular pathway as several genes involved in SUMOylation (Schwartz et al., 2016), however SUMO did not appear to be a component of the aggregates in *Drosophila* Kenyon cells. The possibility remains that SUMOylation of other nuclear proteins may be impacted by HDAC4, however we did not see any alteration in overall pattern of SUMO localization on expression of *3SA*. Given that the deficits in neuronal morphogenesis and memory are facilitated by nuclear HDAC4 which has formed into these aggregates, the identity of other aggregate components and the role of HDAC4 aggregation in neuronal function warrant further investigation.

## Methods

### Fly Strains

All flies were raised on standard medium on a 12-hour light/dark cycle and maintained at a temperature of 25°C unless otherwise indicated. *w[*^*\**^*]; P{w[+mW*.*hs]=GawB}OK107 ey[OK107]/In(4)ci[D], ci[D] pan[ciD] sv[spa-pol]* (*OK107-GAL4*), *w[*^*\**^*]; P{w[+mC]=GAL4-ninaE*.*GMR}12* (*GMR-GAL4*), *P{w[+mW*.*hs]=GawB}elav[c155]* (*elav-GAL4*) and *y1w*^*\**^; *P{UAS-mCD8::GFP*.*L}LL5* (*UAS-CD8::GFP*) were obtained from the Bloomington *Drosophila* Stock Center. *w*^*1118*^; *P{attP,y[+],w[3’] CG42734* (*UAS-MEF2* RNAi) was obtained from the Vienna *Drosophila* Resource Center. *w*^*\**^; *P{w+mC=tubP-GAL80ts}10; TM2/TM6B, Tb1* (*tubP-GAL80ts*) and *w(CS10)* strains were kindly provided by R. Davis (The Scripps Research Institute, Jupiter, FL). FLAG-tagged *Drosophila* and human *HDAC4* and the *3SA* and *L175A* mutant lines have been described previously (Fitzsimons et al., 2013;Schwartz et al., 2016). GFP-tagged *HDAC4* variants were generated by amplification with NotI linkers and insertion of each variant into pUAST-GFP which contained a unique NotI site downstream of GFP to create in frame N-terminal fusions of GFP. *MRE-luc* was generated by annealing the oligos containing 3x consensus MEF2 binding sites (Andres et al., 1995). 3xMREfor 5’ ctagcgatatctgttactaaaaatagaatgttactaaaaatagaatgttactaaaaatagaaa 3’ and 3xMRErev 5’ gatctttc tatttttagtaacattctatttttagtaacattctatttttagtaacagat 3’, digesting with NheI/BglII and inserting into NheI/BglII digested pUAST to create pattBMRE. The minimal hsp70 promoter was PCR amplified from pUASTattB with the following primers hsp70for 5’ atctagatctgagcgccggagtataaatag 3’ and hsp70rev 5’ atcgctcgagggatcccaattccctattcagagttc3’ with BglII and XhoI linkers and cloned into BglII/XhoI of pattB-MRE. pMIR-report (Ambion) was digested with BamHI/SpeI to release the luciferase gene which was cloned into BamHI and XbaI of pattB-MRE to create pattB-MRE-luc. *ΔMRE-luc* was similarly constructed with the oligos 3xΔMREfor 5’ ctagcgatatctgttactaagggtagaatgttactaagggtagaatgttactaagggtaga 3’ and 3xΔMRErev 5’gatctttc tacccttagtaacattctacccttagtaacattctacccttagtaacagatatc 3’. *Drosophila HDAC4* was synthesized by Genscript (New Jersey, USA) (nucleotides 461 – 4216 of NCBI reference sequence NM_132640 with a C-terminal 6x Myc tag) and mutagenesis was performed on this construct to generate the *3SA* and *ΔMEF2* mutants. The *3SA* amino acid substitutions are S239A, S573A and S748A and *ΔMEF2* amino acid substitutions within the MEF2 binding site are K165A L168A and I172A. *DmMEF2* was synthesized by Genscript (nucleotides 1057 – 2601 of NCBI reference sequence NM_057670.5) with a C-terminal 3x HA tag. *DmMEF2* was also similarly subcloned with an N-terminal Myc tag, and the *HDAC4* and *MEF2* constructs were cloned into the pUASTattB plasmid for germline transformation of *Drosophila*. Transgenic flies were generated by Genetivision (Houston, TX) using the P2 docking site at (3L)68A4 for the *HDAC4* constructs, the VK22 docking site at (2R)57F5 for *MEF2-HA*, the (3L)68E1 docking site for *Myc-MEF2* and the VK37 docking site at (2L)22A3 for *MRE-luc, ΔMRE-luc* and *UAS-MEF2-VP16*. All strains were outcrossed for a minimum for five generations to *w(CS10)* flies. Homozygous lines harboring the appropriate GAL4 driver and *tubP-GAL80ts* were generated by standard genetic crosses.

### Scanning electron microscopy

Flies were anaesthetised using FlyNap (Carolina Biologicals) and added to primary modified Karnovsky’s fixative (3% gluteraldehyde, 2% formaldehyde in 0.1 M phosphate buffer, pH 7.2) with Triton X-100 and vacuum infiltrated until soaked through. The flies were then placed in fresh fixative and incubated at RT for at least eight hours. Three 10-minute phosphate buffer (0.1 M, pH 7.2) washes were carried out followed by dehydration using a graded ethanol series for ten to fifteen minutes at each step (25%, 50%, 75%, 95%, 100%) and a final 1-hour wash in 100% ethanol was carried out. Following the ethanol dehydration, the samples were critical point dried using CO_2_ and 100% ethanol (Polaron E3000 series II drying apparatus). The samples were then mounted onto aluminium stubs and sputter coated with gold (Baltex SCD 050 sputter coater) and imaged in the FEI Quanta 200 Environmental Scanning Electron Microscope at an accelerating voltage of 20 kV.

### Behavioral analyses

The repeat training courtship suppression assay (Keleman et al., 2007;Ejima and Griffith, 2011) was used to assess 24-hour long-term memory. Male flies learn that they have been previously rejected by a female and thus when tested with a new female, they display a decrease in courtship behavior compared to naïve males. The detailed methodology has been described previously (Fitzsimons and Scott, 2011;Freymuth and Fitzsimons, 2017). Briefly, single virgin males of each genotype were placed into individual training chambers. A freshly mated wild-type female was placed into half of the chambers (trained group), and then other half of the male group was housed alone (sham group). Flies were incubated in the training chamber for seven hours, during which time multiple bouts of courting were observed in the trained group. The female fly was then aspirated from the training chamber and the males were left in their chambers for the 24 hours prior to testing.

Each trained or naïve male fly was then placed into a testing chamber containing a mated wild-type female and was scored for the time spent performing stereotypic courtship behaviors over the ten-minute period. A courtship index (CI) was calculated as the percentage of the ten-minute period spent in courtship behavior. A mean CI for each group was determined, and from this a memory index (MI) was calculated by the following equation: MI = 1-(mean CI of naïve group/CI of each trained fly) (n≥16/group). The MI was measured on a scale of 0 to 1, a score of 0 indicating memory was no different than untrained naive controls. In all experiments, the scorer was blind to the genotype of the flies.

### Immunohistochemistry

Whole flies were fixed in PFAT/DMSO (4% paraformaldehyde in 1X PBS +0.1% Triton X-100+5% DMSO) for one hour then microdissected in 1xPBS. Brains were post-fixed in PFAT/DMSO for 20 mins and blocked in immunobuffer (5% normal goat serum in 1XPBS+0.5% Triton X-100) for two hours prior to incubation with primary antibody of mouse anti-ELAV, (1:100); mouse anti-FasII (1:200); mouse anti-SUMO-2 (1:20), rabbit anti-GFP (Abcam ab290, 1:1000), rat anti-HA (3F10 Sigma Aldrich, 1:500), mouse anti-Myc (1:100) and rabbit anti-MEF2 (gift from Bruce Paterson, National Cancer Institute, Bethesda, 1:500). They were then incubated with secondary antibody (goat anti-mouse Alexa 488 or 555, goat anti-rabbit Alexa 488 or 555, and goat anti-rat Alexa 555, Sigma Aldrich, 1:500) and mounted with Antifade. The monoclonal antibodies anti-fasciclin II (1D4, developed by G. Goodman), anti-ELAV (9F8A9, developed by G.M. Rubin), anti-Myc (9E 10, developed by J.M. Bishop) and anti-SUMO2 (A82, developed by M. Matunis) were obtained from the Developmental Studies Hybridoma Bank developed under the auspices of the NICHD and maintained by The University of Iowa, Department of Biology, Iowa City, IA 52242. For confocal microscopy, images were captured using a Leica TCS SP5 DM6000B Confocal Microscope and images were processed with Leica Application Suite Advanced Fluorescence (LAS AF) software.

### RT-qPCR

*elav-GAL4* females were crossed to *UAS-MEF2 RNAi* males to generate progeny in which *MEF2* was knocked down in all neurons; and progeny of *elav-GAL4* crossed to *w(CS10)* served as controls. Total RNA was extracted from *Drosophila* heads from three independent crosses for each genotype with the RNeasy Mini kit (Qiagen) according to the manufacturer’s instructions. cDNA was synthesised from 1 ug of total RNA with Transcriptor (Roche) as per the manufacturer’s instructions. RT-qPCR was conducted using SsoFast-EvaGreen (BioRad) reaction master on a Lightcycler II 480 instrument (Roche), following manufacturer’s instructions. The following primers were used: *Mef2*for 5’-GCCACATCACACCCACTCC-3’, *Mef2*rev 5’-GCTGGCCATAGCAGTCGTAG-3’, *EF1a48D*for *5*’*-*ACTTTGTTCGAATCCGTCGC-3’, *EF1a48D*rev 5’-TACGCTTGTCGATACCACCG-3’. A 5-fold dilution of cDNA from control flies was used as template to prepare a standard curve to confirm efficiency of the PCR reactions. Relative quantification was conducted using 2^-ΔΔCt^ method, normalising to the housekeeping gene *Ef1α48D* (Livak et al., 2001).

### Luciferase assays

*elav-GAL4; MRE-luc* and *elav-GAL4; ΔMRE-luc* females were crossed to males of the appropriate genotype. Luciferase assays were performed on the heads of 3-5 day old adults. Heads (10/sample) were homogenized in 20 μL Glo Lysis buffer (Promega), incubated at room temperature for 15 minutes, then centrifuged at 12,000 g for 10 mins. 10 μL of each supernatant was applied per well for each genotype. Quadruplicate samples were analysed for each genotype. 40 μL Bright Glo reagent was added to each well and then luminescence was measured on a PolarStarOMEGA luminometer (BMG Labtech) and normalized to *w(CS10)* control luminescence. Flies were electrically stimulated by placing them in a modified 15 mL plastic tube lined with an electrified grid and exposed to 12 electric shocks (90 V, 1.5 s) over 1 minute and harvested after 1.5 hours to allow transcription and translation of luciferase.

### Transcriptome analysis

Female *elav-GAL4; tubP-GAL80ts* flies were crossed to males individually carrying *UAS-hHDAC4-3SA* or *UAS-hHDAC4-L175A* as well as to *w(CS10)* to provide a control line with the same genetic background and regulatory constructs but no *UAS-HDAC4* construct. Four biological replicates (independent crosses) were generated per genotype. Flies were raised at 18°C then 3-5 day old adult flies were incubated at 30°C for 72 hours to induce expression of *3SA* and *L175A* in all neurons. The flies were then snap frozen in liquid nitrogen and their heads separated by vortexing and sorted from bodies by sieving. RNA extracted from *Drosophila* heads (RNeasy mini kit, Qiagen) was quantified on a Denovix DS-11 spectrophotometer and assessed for quality on a Labchip GX Touch HT (PerkinElmer). Illumina HiSeq 150 bp paired-end sequencing was carried out by Novogene (Hong Kong). The sequencing yielded an average paired end read count of 36.9 million per sample replicate. Read mapping and read count analysis was performed by NextGen Bioinformatic Services (New Zealand). The raw read data was adaptor filtered and quality trimmed (phred>10) using BBDuk (version 36.86; BBTools by B Bushnell www.sourceforge.net/projects/bbmap/). Following quality trimming any reads shorter than 50 bp were removed to minimise the possibility of multiple mapping. The BBDuk parameters were: minlen=50 qtrim=rl trimq=10 ktrim=r k=21 mink=11. The data was observed to be high quality with over 99% of bases passing the filtering and quality trimming step. High quality reads were mapped to the *Drosophila* genome (release 92) downloaded from ENSEMBL using HISAT2 (version 2.05) in stranded mapping mode. The alignment rate to the genome was >94% of reads for all sample replicates. RNA-seq reads from the *3SA* and *L175A* libraries were also mapped to human chromosome 2 (assembly version GRCh38) to verify the presence of the mutations in the samples from of these two treatment groups. Gene based read counts were generated from the alignment BAM files using HT-Seq (version 0.6.0) based on the ‘union’ mode. The count files were processed in R (version 3.4.0) and differentially expressed genes were identified using the package DESeq2 (version 1.18.1) as described in the vignette (Love et al., 2014). Differentially expressed genes with an FDR < 0.05 (i.e. significantly altered) were uploaded to the Database for Analysis, Visualization and Integrated Discovery (DAVID, 6.8)(Huang da et al., 2009) to identify functional groups highly enriched in a sample, as well as identifying biological pathways that are highly represented in a given group.

**Table S1. Differentially expressed genes between control and L175A; control and 3SA; and 3SA and L175A groups**. Genes with a False Discovery Rate of <0.05 are shown.

## Acknowledgements

We thank Sangeeta Chawla for the MEF2-VP16 plasmid (University of York) and Bruce Paterson for the anti-MEF2 antibody (National Cancer Institute, Bethesda). We also thank the Manawatu Microscopy and Imaging Centre, Massey University for assistance with confocal and SEM.

## Author Contributions

HF, PM and WTJ designed the study. HF directed the study and carried out the luciferase and anti-SUMO IHC experiments. PM carried out the majority of the experiments. WJT performed light microscopy, SEM and immunohistochemistry on the double mutant. DW performed the bioinformatic analyses. HF wrote the manuscript with assistance from PM and WJT.

## Data Availability Statement

The NCBI bioproject number for this study is PRJNA664892 (https://www.ncbi.nlm.nih.gov/bioproject/PRJNA664892) and the raw sequencing data is available at the NCBI Short Read Archive (https://www.ncbi.nlm.nih.gov/sra) under accession numbers SRR12689988-SRR12689999

