## Supplemental Figure 1 for "Increased abundance of nuclear HDAC4 impairs neuronal development and long-term memory"

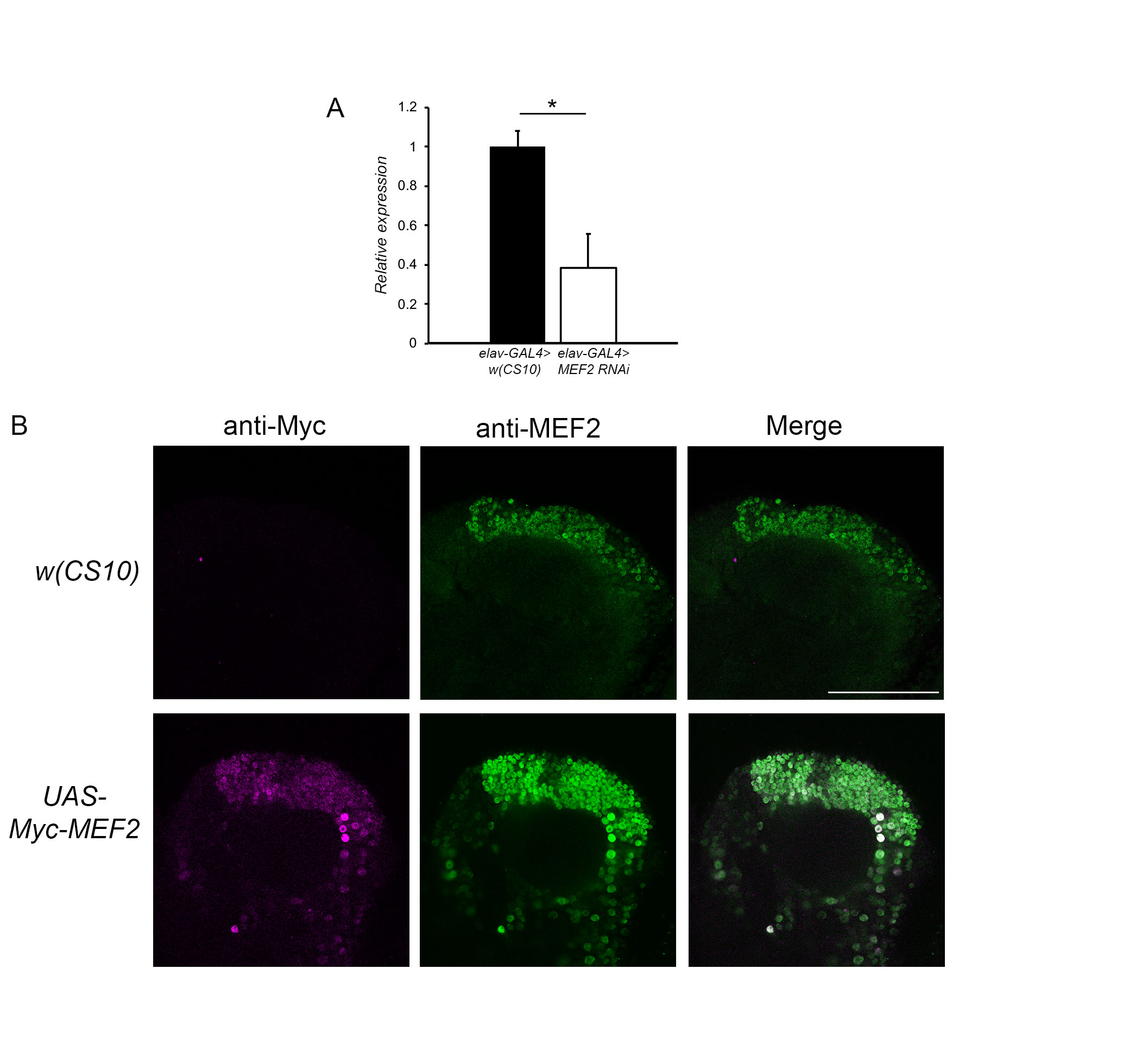

**Figure S1. Confirmation of MEF2 knockdown and overexpression.** A. *elav-GAL4* females were crossed *UAS-MEF2 RNAi* males in order to knockdown *MEF2* in all neurons. RNA was isolated from heads and RT-qPCR for *MEF2* was carried out with normalization to *Ef1α48D. MEF2* was significantly knocked down to approximately 40% of wild-type *(t*-test t_(4)_=3.99, p<0.05)*.* B*. OK107-GAL4; tubP-GAL80*_ts_ females were crossed to *UAS-Myc-MEF2* or *w(CS10)* males. Flies were raised at 18°C until after eclosion, then placed at 30°C 72 hours prior to dissection to induce transgene expression. B. Whole-mount brains were subjected to immunohistochemistry with anti-MEF2 (green) and anti-Myc (magenta). Optical sections (1 μm) through the calyx of the mushroom body shows that Myc-MEF2 co-localises with endogenous MEF2 in Kenyon cell nuclei, whereas there is no Myc-MEF2 expression in the control. Scale bar = 50 μm.
